# Functional characterization of *Salmonella* Typhimurium encoded YciF, a domain of unknown function (DUF892) family protein, and its role in protection during bile and oxidative stress

**DOI:** 10.1101/2023.01.20.524870

**Authors:** Madhulika Singh, Aravind Penmatsa, Dipankar Nandi

**Affiliations:** Department of Biochemistry, Indian Institute of Science, Bangalore-560012, India; Molecular Biophysics Unit, Indian Institute of Science, Bangalore-560012, India

## Abstract

YciF (STM14_2092) is a member of domain of unknown function (DUF892) family. It is an uncharacterized protein, involved in stress responses in *Salmonella* Typhimurium. In this study, we investigated the significance of YciF and its DUF892 domain during bile and oxidative stress responses of *S*. Typhimurium. Purified wild type YciF forms higher order oligomers, binds to iron and displays ferroxidase activity. Studies on the site-specific mutants revealed that the ferroxidase activity of YciF is dependent on the two metal binding sites present within the DUF892 domain. Transcriptional analysis displayed that the Δ*csp*E strain, which has compromised expression of YciF, encounters iron toxicity due to dysregulation of iron homeostasis in presence of bile. Utilizing this observation, we demonstrate that the bile mediated iron toxicity in Δ*csp*E causes lethality, primarily through the generation of reactive oxygen species (ROS). Expression of wild type YciF, but not the three mutants of the DUF892 domain, in Δ*csp*E alleviate ROS in presence of bile. Our results establish the role of YciF as a ferroxidase that can sequester excess iron in the cellular milieu to counter ROS-associated cell death. In fact, pre-treatment with an iron chelator mitigates the hypersensitivity of Δ*csp*E to bile.

**Importance:** The DUF892 domain has a wide taxonomic distribution encompassing several bacterial pathogens. This domain belongs to the ferritin-like superfamily; however, it has not been biochemically and functionally characterized. This is the first report of characterization of a member of this family. In this study, we demonstrate that *S*. Typhimurium YciF is an iron binding protein with ferroxidase activity, which is dependent on the metal binding sites present within the DUF892 domain. It combats iron toxicity and oxidative damage caused due to exposure to bile. The functional characterization of YciF delineates the significance of the DUF892 domain in bacteria. In addition, our studies on *S*. Typhimurium bile stress response divulged the importance of comprehensive iron homeostasis in bacteria in presence of bactericidal compounds that tend to generate ROS irrespective of their primary targets.

## Introduction

*Salmonella enterica* serovar Typhimurium (*S*. Typhimurium) is an enteric pathogen that causes non-typhoidal Salmonellosis or gastroenteritis. The global burden of gastroenteritis caused by non-typhoidal *Salmonella* (NTS) is substantial (1). Besides, *S*. Typhimurium can also be invasive to usually sterile sites such as blood and cause bacteraemia in immunocompromised individuals (2–4). The increasing antibiotic resistance in *S*. Typhimurium has resulted in formation of multiple drug resistant strains and presents a significant global health challenge (5). Infection and survival within host is aided by myriad of stress response genes responsive to various environmental stressors. Bile is one such stressor with antibacterial activity. As a pathogen *Salmonella enterica* manifests an extreme example of bile resistance because it can colonize the hepatobiliary tract during systemic infection and the gall bladder during chronic infection (6). Bile is a potent antimicrobial agent that causes cell membrane damage, protein denaturation, lipid peroxidation, secondary structure formation in RNA, DNA damage. In addition, it can also chelate important minerals such as calcium and iron (6, 7). Studies have reported that bile can lead to oxidative stress within the cells (8, 9). Although the RpoS mediated general stress response pathway is critical to survival of *S*. Typhimurium in bile, an RpoS-independent pathway also exists which is mediated by CspE (6, 10). CspE belongs to the family of cold shock proteins and has significance in diverse stress response pathways as it an RNA chaperone that stabilizes the transcripts of several stress response proteins (11). In *S*. Typhimurium 14028s it is critical for tolerance to bile stress and deletion of this gene leads to hypersensitivity to bile (10).

Protein families are annotated as ‘domain of unknown function’ (DUF) when they have no member with an experimentally determined function. DUF892 is functionally uncharacterized and belongs to ferritin-like superfamily (12). Ferritins maintain the supply of iron to the cells. Besides iron storage, ferritins help in protection against free radicals generated due to Fenton reaction when there is iron excess (13, 14). For *Salmonella*, iron is a crucial resource however its bioavailability is a significant challenge and excess is hazardous (15). *S*. Typhimurium encounter an iron restricted environment inside the host as a result of nutritional immunity utilized by the host (15, 16). In response, the bacteria also have evolved effective iron acquisition approaches such as production of siderophores to capture iron (16, 17). While the problem associated with ferric ion is that it has low solubility in physiological conditions, ferrous ions can cause cytotoxicity. Excess ferrous ions activate dioxygen leading to formation of intermediate reactive species that results in generation of ROS and damage proteins and DNA (18, 19, 20). Therefore, it is important for cells to strictly regulate the intracellular iron content.

YciF is an all α-helical protein containing a single domain DUF892. As a member of domain of unknown function (DUF892) family, YciF is functionally uncharacterized, as is the DUF892 domain. It contains two metal binding sites as revealed by the crystal structure (21, 22). In *S*. Typhimurium there have been few studies that suggest YciF is a stress response protein. YciF has been found to be highly upregulated in osmotic shock and bile stress (10, 23, 24). In bile stress it is shown to work downstream of CspE (10). Although, YciF has been described as a bacterial stress response protein, the biochemical function as well as the intracellular mechanism of action is unknown.

Previously, our lab reported that overexpression of the uncharacterized protein, YciF confers survival advantage to bile sensitive *S*. Typhimurium cspE deletion strain (Δ*csp*E) (10). We chose to investigate the significance of YciF and its DUF892 domain in the bile stress response of *S*. Typhimurium. In this study, YciF and mutants of the metal binding site present in DUF892 domain were purified, characterized biochemically and physiologically. We found that YciF has ferroxidase activity which is compromised when the metal binding site is mutated. To associate the biochemical activity of YciF with the *in vivo* phenotype, we determined the presence of iron toxicity in the bile treated Δ*csp*E strain. Furthermore, we show that the high intracellular ROS level in the Δ*csp*E strain, possibly resulting from bile mediated iron dysregulation, is lowered in presence of YciF. YciF was also observed to directly rescue the peroxide sensitive phenotype of the Δ*csp*E strain, highlighting its role in mitigating deleterious effects of ROS.

## Results

### YciF contains diiron sites within the DUF892 domain

*S*. Typhimurium encoded YciF has a single ferritin-like domain, DUF892 (Fig. 1A). However, the functional significance of this domain remains obscure as members of the DUF892 family are uncharacterized. The DUF892 domain homologs are found across members of Gammaproteobacteria, Betaproteobacteria, Alphaproteobacteria, Bacteroidetes and even Archaeabacteria (Fig. S1). *S*. Typhimurium YciF shows high sequence identity with other members of Enterobacteriaceae (>80%) (Fig. 1B). The DUF892 domain of YciF comprises two metal binding sites identical to diiron centre. The putative metal binding sites are more conserved among members of Enterobacterales (Fig. S1). The diiron centre containing proteins are of several types, characterized by the presence of glutamate and histidine as part of ExxH motif and two additional glutamates inside a four-helix bundle (12). Diiron proteins such as ribonucleotide reductases, bacterioferritin have two carboxylates as bridging ligand whereas ferritins have a single carboxylate ligand for the diiron sites (12, 25). Similar to ferritins, YciF has E143D as the single carboxylate bridging ligand (Fig. 1C).

**Fig 1.**
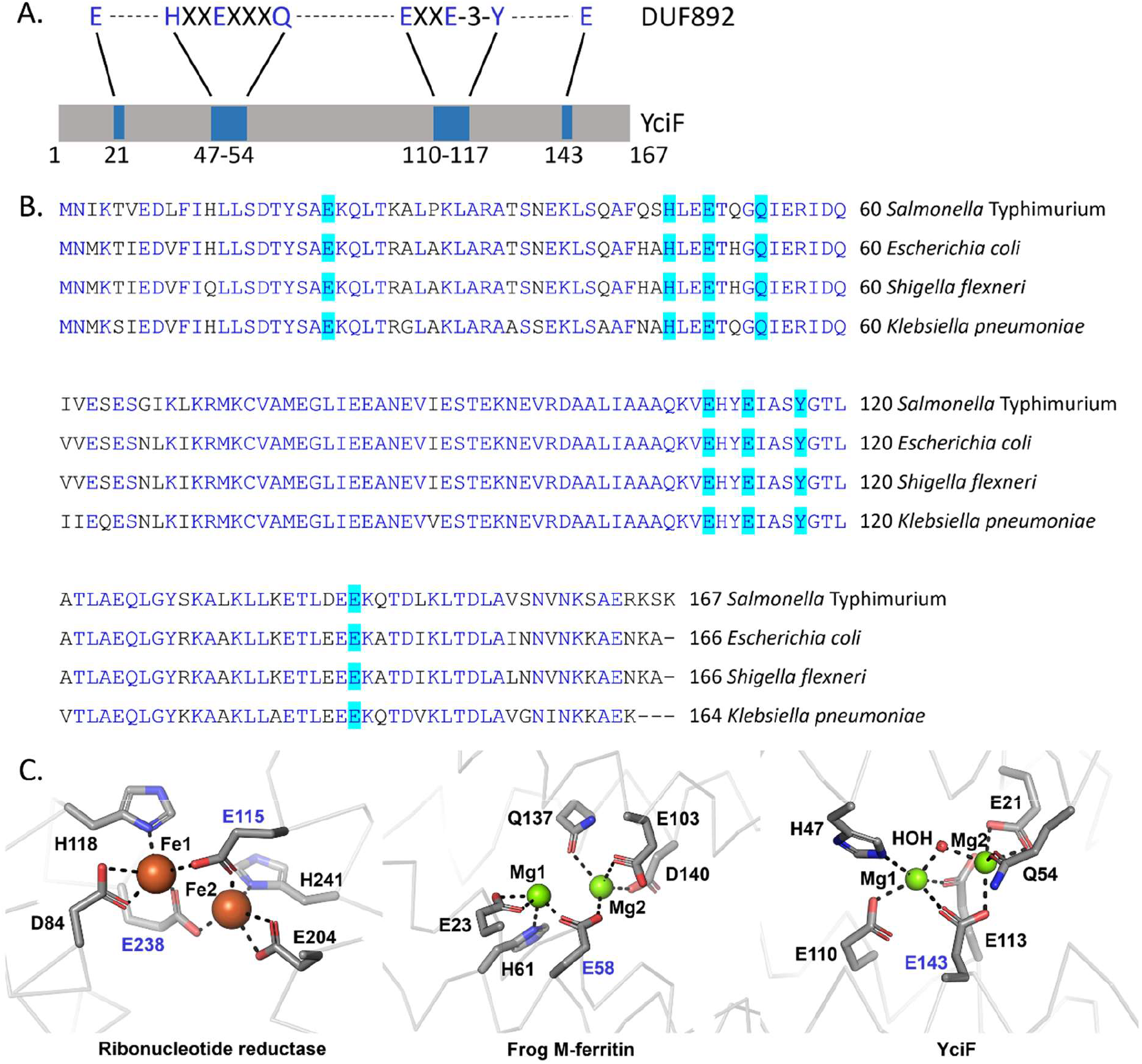
(A) YciF contains a single ferritin-like domain, DUF892 that spans almost the entire length of the protein. (B) Multiple sequence alignment using clustal omega shows that YciF is highly conserved across Enterobacteriaceae. The residues of DUF892 domain involved in metal ion-coordination are highlighted in turquoise. (C) Comparison of diiron centres of ribonucleotide reductase (1PFR), frog M-ferritin (1MFR) and YciF (4ERU). E143 and E58 are the single bidentate residues for YciF and frog M-ferritin, respectively whereas ribonucleotide reductase has two carboxylate bridging residues (E115 and E238).

### YciF and its metal binding sites mutants form higher order oligomeric complex

*S*. Typhimurium YciF crystal structure has two metal binding sites (22). To understand the role of DUF892 domain that contains the expected diiron site, single point mutants Q54A and E113Q were generated targeting the two metal binding sites, M2 and M1 respectively. A third mutant E143D was made that would disrupt coordination of both the metal binding sites as E143 is the bridging ligand according to the YciF crystal structure (Fig S2). YciF and the mutants (Q54A, E113Q, E143D) were purified using the streptactin affinity resin and subjected to SDS PAGE (Fig. 2A). The stability of the mutants was determined using thermal shift assay. E113Q and Q54A had almost similar melting temperature as YciF whereas E143D had decreased melting temperature. However, E143D was stable enough as the Tm of 53.5°C is still substantially higher than the physiological temperature of 37°C (Fig. S3). Members of the ferritin superfamily are known to form higher order oligomeric complexes (13, 26, 27). To investigate the possibility of YciF forming higher order oligomer, gradient Native PAGE was performed. YciF and the mutants had electrophoretic mobility corresponding to that of 150 kDa Y-globulin (Fig. 2B). YciF was subjected to SECMALS to determine the precise molecular weight of the oligomeric complex. It assumes a 120 kDa complex under native conditions (Fig. S4). Using the SEC-MALS data as reference, analytical size exclusion chromatography was performed to compare the oligomeric status of YciF and its mutants. All three mutants formed oligomeric complexes that are comparable to the unmodified YciF (Fig. 2C). The approximate molecular weight of the proteins was calculated using protein with standard masses. The SEC data correlates with the SEC-MALS analyses wherein WT YciF oligomer had an estimated molecular mass of 109 kDa (±3 kDa). Q54A, E113Q and E143D formed oligomer corresponding to 104, 100 and 103 kDa respectively (Fig. 2D).

**Fig 2.**
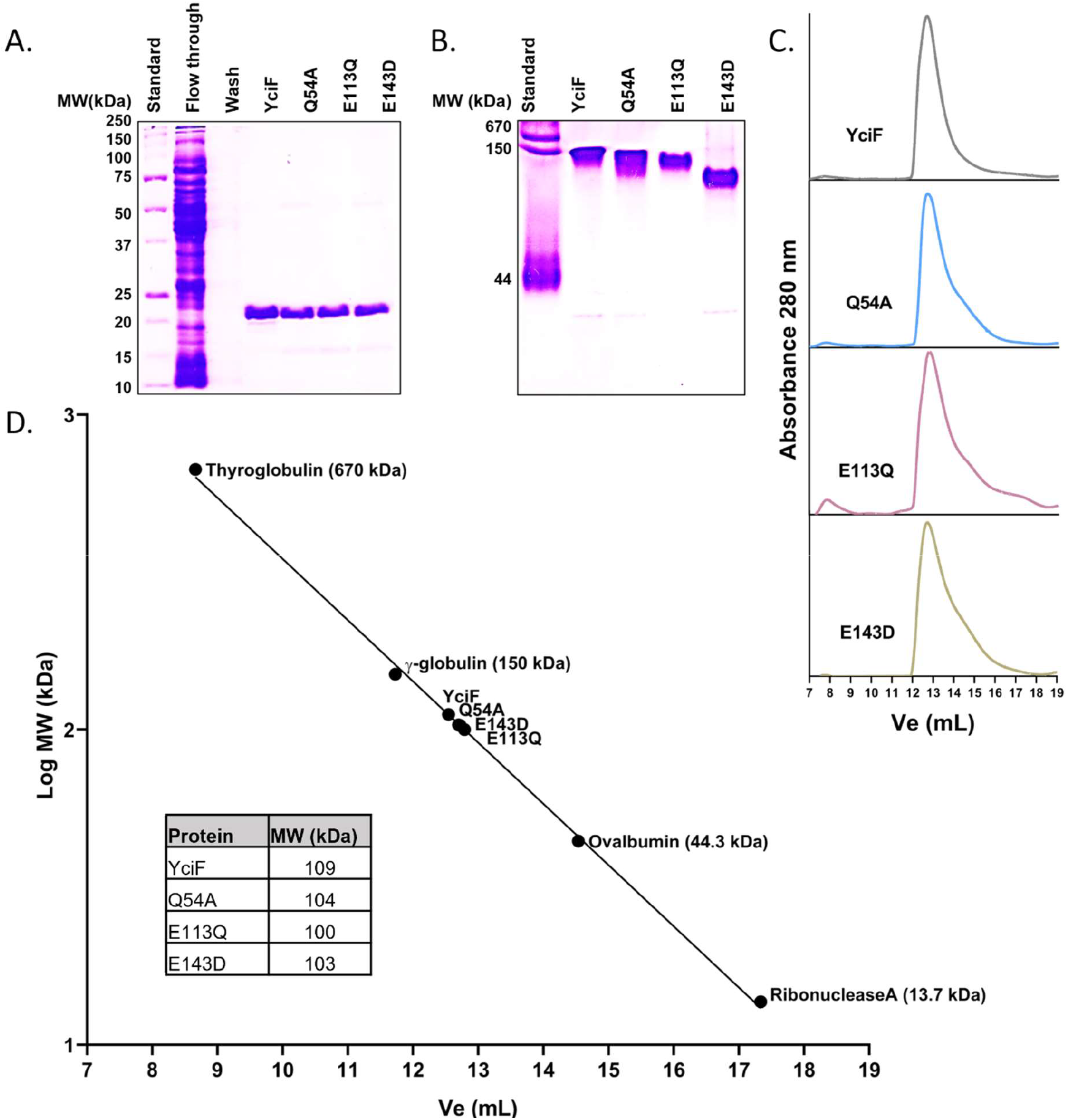
*S*. Typhimurium YciF and its metal binding sites mutants form higher order oligomers. (A) 12.5% SDS PAGE profile of Strep II tagged YciF, Q54A, E113Q and E143D. Monomeric YciF and its mutants have a molecular weight of approximately 20 kDa. (B) 4-15% gradient native PAGE profile of YciF, Q54A, E113Q and E143D. (C) Analytical size exclusion chromatography to determine the apparent molecular weight of YciF and its mutants. Purified proteins were subjected to Superdex S-200 column. Molecular weight standard was run on the same column maintaining similar elution conditions. The Table inside represents the molecular mass calculated from linear regression equation.

### YciF interacts with iron and exhibits ferroxidase activity

The domain architecture of YciF suggests that it could bind iron. However, in the crystal structure magnesium ion occupies the metal binding sites (22). Therefore, thermal shift assay was performed to identify the native ligand of the protein. In the presence of Mg^2+^, there was a moderate increase in melting temperature while addition of Zn^2+^ slightly destabilized the protein, resulting in reduced melting temperature (Fig. 3A). A distinctive melt curve was obtained in presence of iron where no derivative peak was observed till 95°C suggesting that melting temperature could be higher than 95°C as has been observed in case of ferritin (28–31) or the protein was stable till aggregating temperature. No significant alteration was observed in melting temperature in presence of metal chelator EDTA indicating that metal ion is unlikely to be a cofactor for the protein (Fig. 3A). For comparison of iron incorporation into YciF, Q54A, E113Q and E143D, protein samples were incubated with iron and subjected to Native PAGE. Gel was stained with Ferene-S staining solution to detect iron bound to protein. At sub-saturating concentration of iron, YciF was able to bind and retain the iron whereas the mutants did not (Fig. 3B).

**Fig 3.**
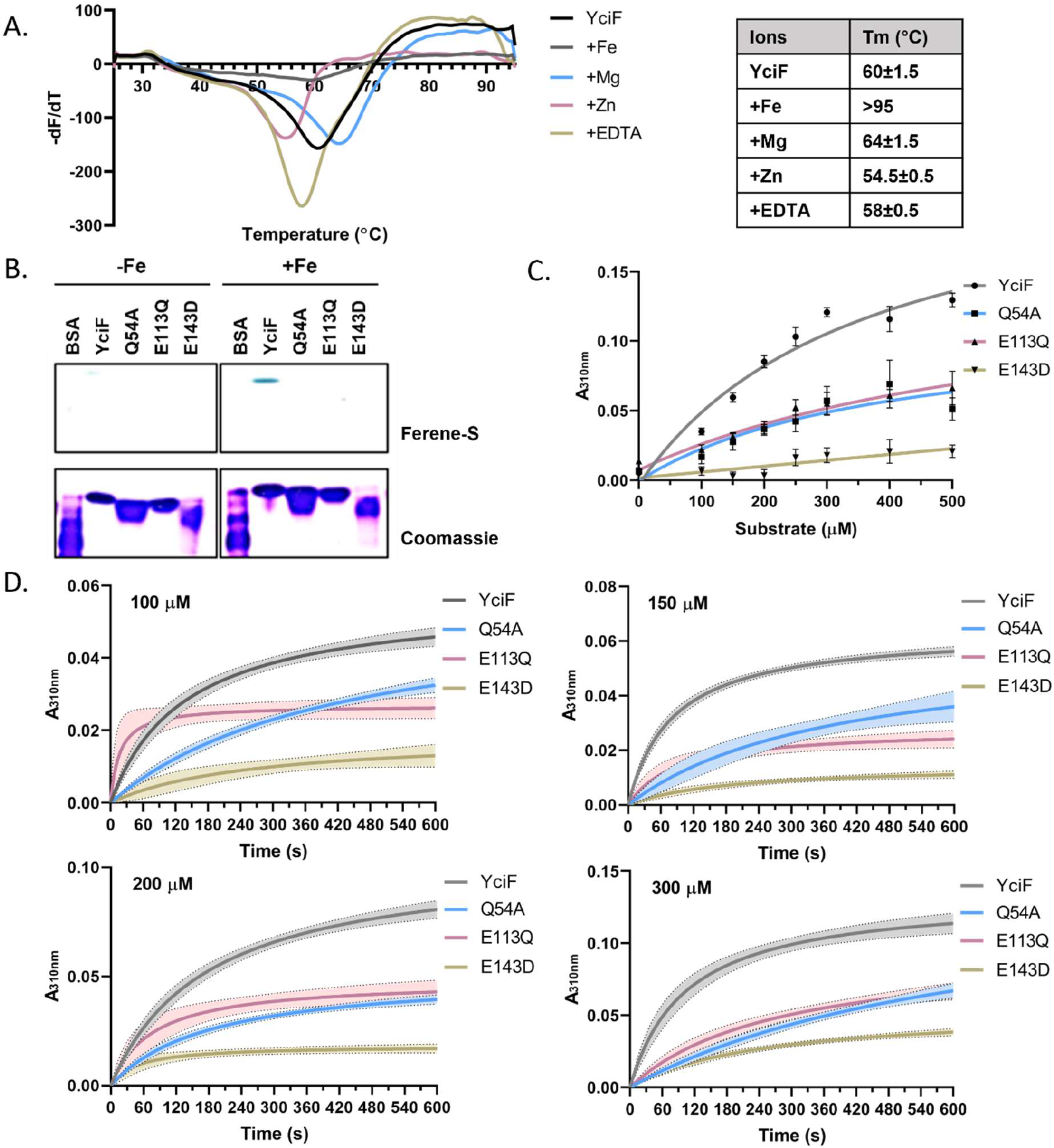
*S*. Typhimurium YciF binds to iron and displays ferroxidase activity. (A) Thermal shift assay was performed to determine the native ligand. 10 pg YciF was incubated with 200 μM (NH_4_)_2_Fe(SO_4_)_2_, MgSO4, ZnCl2 and EDTA. The table represents the melting temperature (Tm). (B) Comparison of iron binding of YciF and its metal binding sites mutants. Purified proteins were incubated with 200 μM (NH_4_)_2_Fe(SO_4_)_2_ and subjected to native PAGE followed by staining with Ferene-S and Coomassie brilliant blue. BSA was used as negative control. (C) Determination of ferroxidase activity of YciF and the mutants at different concentrations of (NH_4_)_2_Fe(SO_4_)_2_). Formation of ferric ion was measured at 310 nm. (D) Kinetic comparison of ferroxidase activity of YciF, Q54A, E113Q, E143D. (NH_4_)_2_Fe(SO_4_)_2_) was added at different concentrations and absorbance at 310 nm was measured for 600 s at 30 s interval. Values were blanked against no protein control to correct for auto-oxidation of ferrous ion. Graphs were plotted using non-linear regression. The data is representative of 3 independent protein preparations.

As the diiron center of YciF is similar to ferritin, we decided to study the ferroxidase activity. The ferroxidase center catalyzes ferrous ions and releases the ferric products which can be measured at absorbance wavelength of 310 nm. Initial optimization of conditions for ferroxidase assay was performed at different pH, buffers, and protein concentrations (Fig. S5A, B, C). Oxidation of ferrous ion to ferric ion was faster in presence of YciF than the auto-oxidation of ferrous ion. Oxidation of different concentrations of Mohr’s salt by YciF was monitored aerobically at 310 nm. Q54A and E113Q with mutations in the M2 and M1 sites respectively, had compromised activity compared to YciF while E143D that had mutation in the bridging amino acid did not display any significant ferroxidase activity (Fig. 3C). Ferroxidase reaction progression was measured kinetically at 30s intervals wherein the result corroborated with the substrate titration data (Fig. 3D). A colorimetric method that utilizes Ferene-S as a ferrous ion-specific chromogen was also used to determine the ferroxidase activity of YciF. Loss of ferrous ion due to catalytic activity of YciF led to decrease in absorbance at 590 nm. Notably, all the three mutants displayed reduced activity (Fig. S5D).

### The Δ*csp*E strain shows dysregulation of iron homeostasis genes upon bile treatment

Previously, our laboratory had reported that the bile sensitive phenotype of Δ*csp*E could be rescued by overexpressing YciF, however the mechanisms were not studied (10). Therefore, we wanted to elucidate the underlying mechanism of function of the ferroxidase activity of YciF during bile stress response. As ferroxidase activity is known to be primarily associated with mitigating iron toxicity within bacterial cells, we investigated whether there was any iron imbalance in the Δ*csp*E strain possibly resulting from bile treatment. For this, transcript levels of three different categories of iron homeostasis genes involved in uptake of iron, intracellular release of iron and iron storage were analyzed. *fep*A is required for uptake of ferri-enterobactin complex into the cell while ferrous ion can directly pass through *feo*B and *sit*A. *fhu*C is a ferri-hydroxamate transporter and its upregulation suggests that bile could lead to iron starvation or act as a cue for the pathogen of an iron-deprived environment such as host gut. *S*. Typhimurium is mainly known to produce enterobactin and salmochelin type of siderophores (32, 33) whereas hydroxamate siderophores are produced by other microbes in gut but can be utilized by *Salmonella* through its hydroxamate transporters (34, 35). All the genes involved in uptake of iron were upregulated in bile stress. However, the upregulation of *fep*A, *fhu*C and *feo*B was significantly higher whereas sitA also showed increased level of transcripts in bile treated Δ*csp*E compared to the WT strain (Fig. 4A). To understand whether the iron is released from the ferri-siderophore complex into the cytoplasm, transcript levels of fes (ferri-enterobactin utilization), fhuF (ferrioxamine utilization) and *yqj*H (ferric reductase) were analyzed. Upregulation of these genes in *S*. Typhimurium WT upon bile treatment suggests that the iron acquired through siderophores is released into the cytoplasm. Δ*csp*E had higher expression of these genes than the WT strain which was significant (Fig. 4B). *ftn*B and *dps* encode proteins that are involved in storage of iron and provide protection against iron-dependent oxidative stress mediated killing (11). Downregulation of both the genes was observed in Δ*csp*E in untreated condition. A decrease in transcript levels of these genes was observed following bile treatment in WT which substantially reduced further in Δ*csp*E. Fur is a negative regulator of iron uptake in cells but no change was observed in its transcript levels (Fig. 4C). These results indicate that possibly there is a considerable surge in intracellular iron content of Δ*csp*E compared to the WT strain during bile stress.

**Fig 4.**
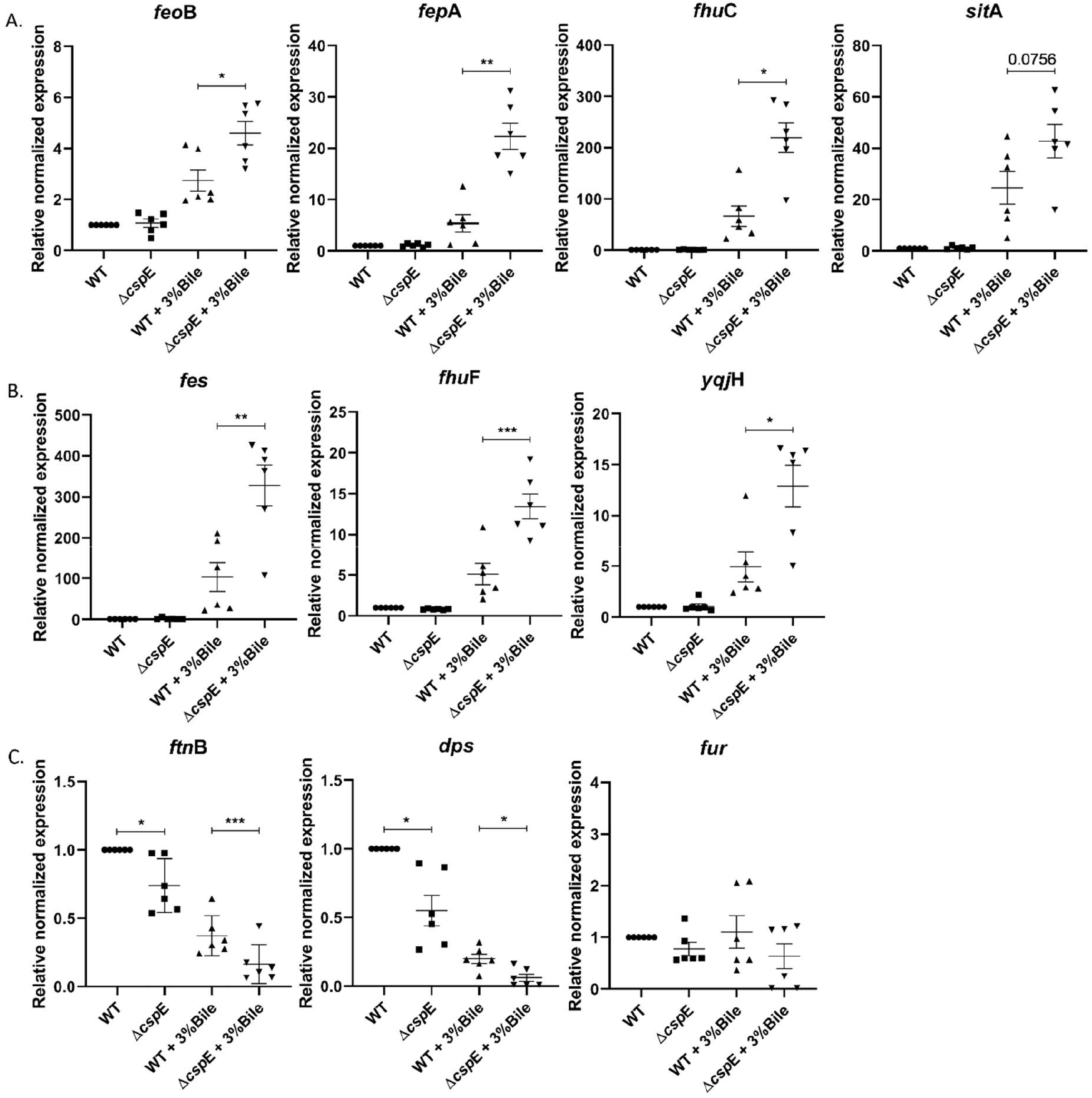
qRT-PCR analysis of genes responsible for iron balance in *S*. Typhimurium reveals iron disbalance in Δ*csp*E in presence of bile. (A) Genes encoding ferri-siderophore transporters: *fep*A, *fhu*C are responsible for uptake of Fe^3+^ while *sit*A and *feo*B are involved in Fe^2+^ uptake. (B) Genes encoding enzymes responsible for intracellular release of iron from siderophores. (C) Genes regulating intracellular iron level: *ftn*B and dps are iron storage proteins whereas fur is a negative regulator of iron uptake. Cq value of WT strain grown in LB was used to normalize Cq values of all the panels. Data shown as mean±SEM and is representative of 6 independent experiments. P values were measured by one-way ANOVA using Sidak’s multiple comparisons test. *p<0.05, **p<0.01, ***p<0.001.

### The Δ*csp*E strain has higher intracellular ROS compared to WT following bile treatment which reduces significantly upon YciF overexpression

Disruption of intracellular iron homeostasis can lead to production of reactive oxygen species. Bile treatment is also known to cause oxidative stress inside bacterial cells. As Δ*csp*E showed iron dysregulation at the level of transcription, we investigated whether Δ*csp*E has increased ROS levels, possibly resulting from bile treatment coupled with higher intracellular free iron content. In addition, we studied the effects of overexpression of YciF which shows ferroxidase activity, in changing intracellular ROS levels. Upon bile treatment, Δ*csp*E strain containing vector alone indeed showed enhanced ROS generation compared to WT strain containing vector alone. Overexpression of YciF led to decrease in intracellular ROS in Δ*csp*E and there was significant reduction in DCFDA positive cells compared to vector control. WT strain containing vector alone or p*yci*F had similar levels of ROS upon bile treatment (Fig. 5A and B). Total ROS levels were compared between bile treated cells overexpressing YciF and the mutants. The Δ*csp*E strain overexpressing Q54A, E113Q and E143D showed reduction in ROS levels compared to vector control. Nevertheless, in these strains ROS was still higher compared to YciF overexpression condition (Fig 5C). We observed that production of ROS was accompanied by cell lethality. Phenotypically, Δ*csp*E strain overexpressing Q54A, E113Q and E143D mutants were still susceptible to bile stress although YciF significantly rescued the bile sensitive Δ*csp*E strain.

**Fig 5.**
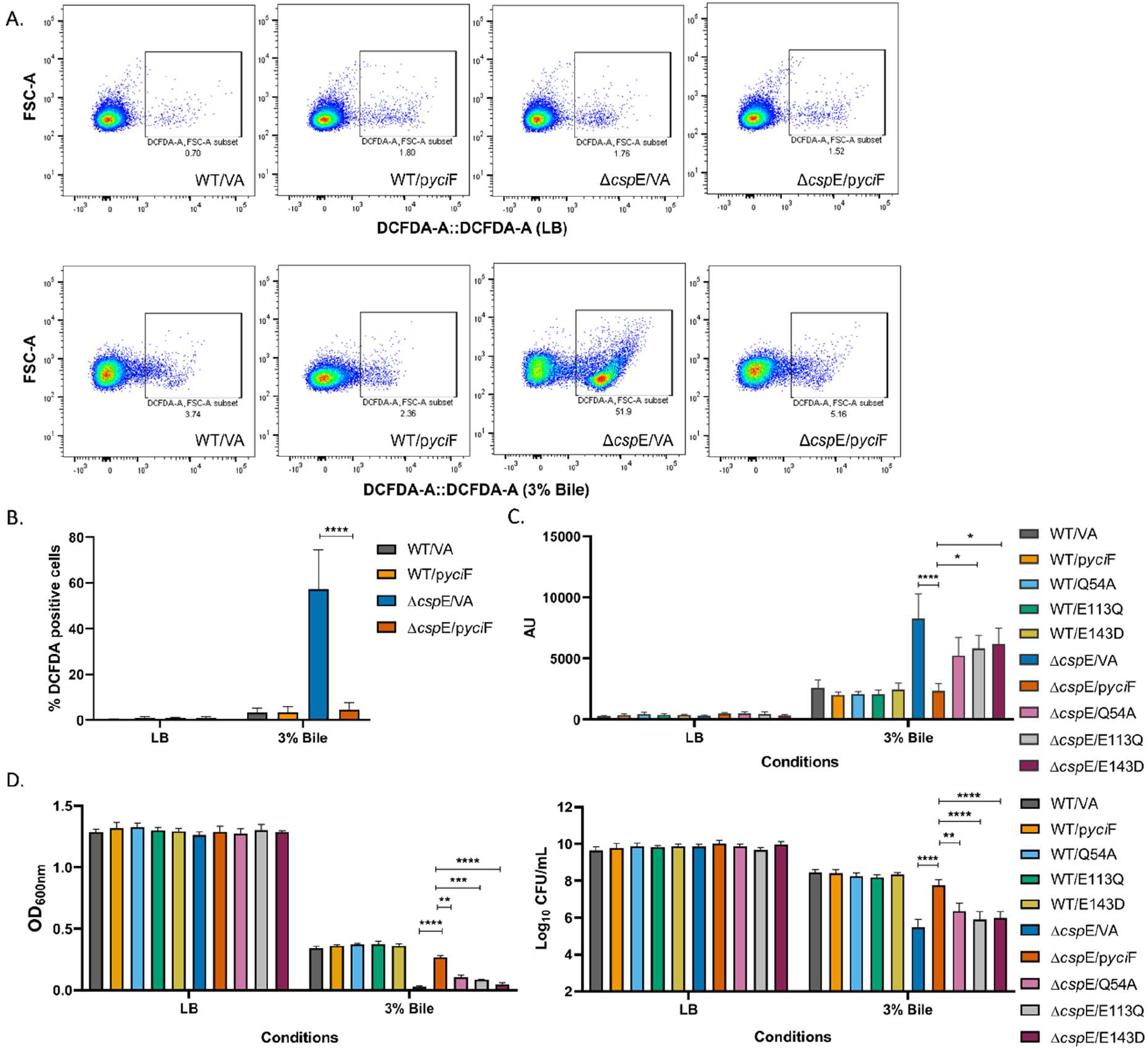
YciF combats bile mediated oxidative stress. (A) S.Typhimurium WT and Δ*csp*E strains overexpressing YciF were grown in absence or presence of 3% bile for 6 hours. Cells were stained with DCFDA and analysed by flow cytometry. (B) The percentage DCFDA positive cells based on flow cytometry data. (C) Quantitation of intracellular ROS levels upon 3% bile treatment in strains overexpressing YciF and the metal binding sites mutants using DCFDA staining. Fluorescence was measured using excitation and emission wavelength of 485 nm and 535 nm respectively. (D) Growth of strains overexpressing YciF and the mutants in absence or presence of 3% bile determined by measuring O.D. at 600nm. Cells were plated on LB agar post bile treatment and colonies were counted. Data shown as mean±SEM and is representative of 4 independent experiments. P values were analysed by two-way ANOVA using Sidak’s multiple comparisons test. *p<0.05, **p<0.01, ***p<0.001, ****p<0.0001.

### Chelation of iron increases the survival of the Δ*csp*E strain upon bile treatment

As Δ*csp*E showed imbalance of transcripts involved in iron homeostasis we assessed any growth differences between *S*. Typhimurium WT and Δ*csp*E strains in presence of the iron chelator 2,2 bipyridyl. Both the strains had similar growth across different doses of 2,2 bipyridyl (Fig. S6). High levels of intracellular iron and ROS content as a result of bile stress can lead to Fenton reaction and thereby production of hydroxyl radicals that is toxic to the cell (36). CspE is known to be critical for survival of *S*. Typhimurium 14028s strain in bile stress condition and the Δ*csp*E strain was highly susceptible to bile mediated killing compared to the WT strain, in agreement to the previous study (10). The WT strain had similar growth in bile irrespective of the absence or presence of 2,2 bipyridyl. However, the severe growth attenuation of Δ*csp*E observed upon bile treatment was rescued when media was supplemented with 2,2 bipyridyl (Fig. 6A and B). The increase in growth of Δ*csp*E in presence of bile upon iron chelation was accompanied with reduction in ROS levels. No alteration was observed in the ROS levels of bile treated WT cells in presence of 2, 2 bipyridyl. (Fig. 6C). These results clearly demonstrate that the Δ*csp*E strain is susceptible to iron mediated killing in presence of bile and can grow similar to WT strain if excess free iron is sequestered away. Furthermore, the Δ*csp*E strain was found to be sensitive to H_2_O_2_ compared to the WT and the growth inhibition was overcome when YciF was overexpressed (Fig. 6D).

**Fig 6.**
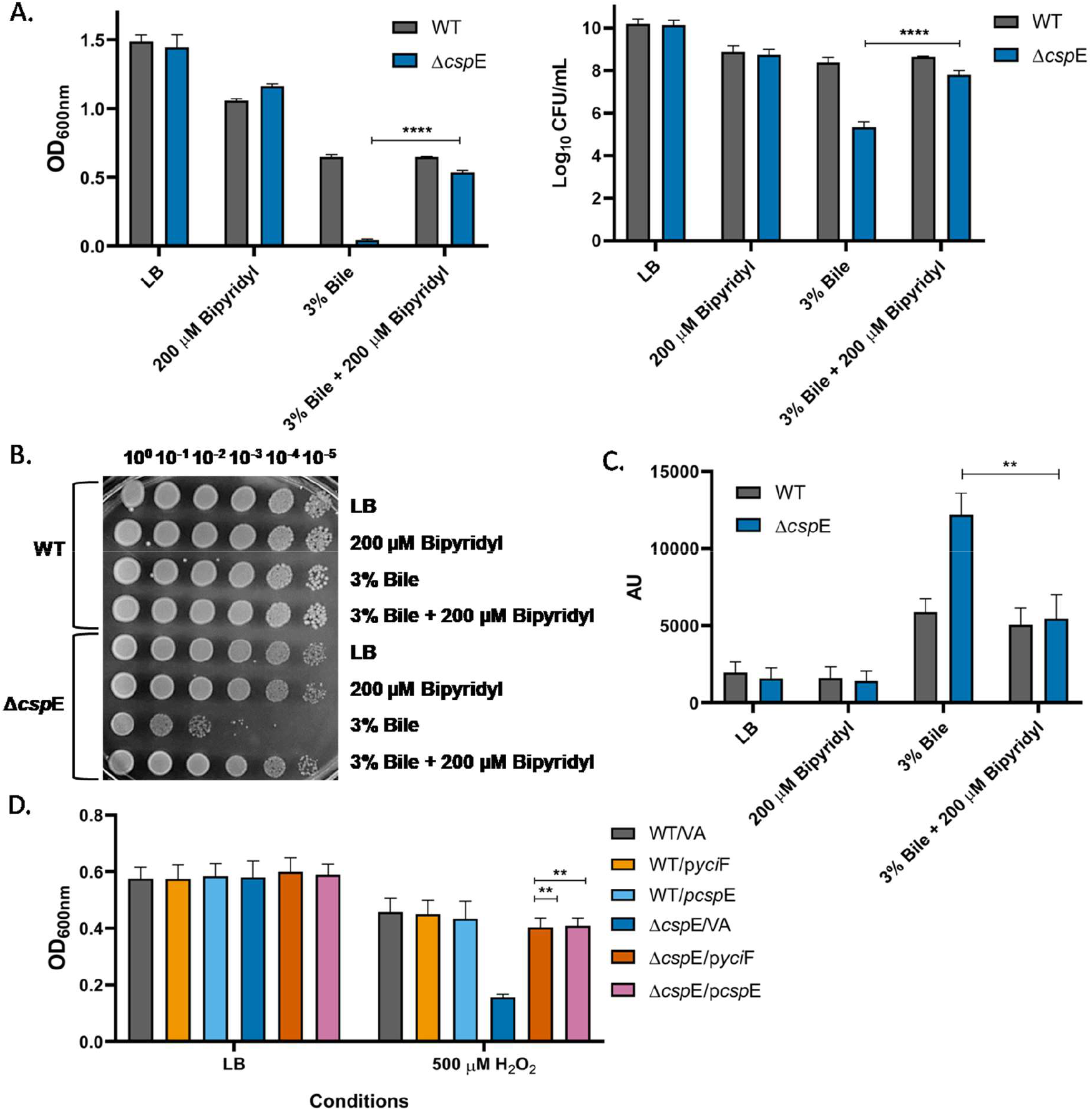
Iron chelation reduces ROS and increases the growth of Δ*csp*E strain in presence of bile. S.Typhimurium WT andΔ*csp*E were grown in LB alone and LB containing 200 μM bipyridyl/ 3% bile/ 3% bile + 200 μM bipyridyl for 6 hours. For H_2_O_2_ treatment, the mentioned strains were grown for 3 hours. (A) O.D. was measured at 600 nm and cells were plated on LB agar following 3% bile treatment and colonies were counted. (B) WT and Δ*csp*E strains were spotted onto LB agar plates after 6 hours of 3% bile treatment. (C) Cellular ROS levels in absence or presence of 2,2-bipyridyl following 3% bile treatment was quantitated using DCFDA staining. Fluorescence was determined at excitation and emission wavelength of 485 nm and 535 nm respectively. (D) O.D. was measured at 600 nm after H_2_O_2_ treatment. Data shown as mean±SEM and is representative of 4 independent experiments. P values were measured by two-way ANOVA using Sidak’s multiple comparisons test. **p<0.01, ****p<0.0001.

## Discussion

Previous studies have shown that CspE is important for bile tolerance as well as biofilm formation in *S*. Typhimurium (10, 37). Physiologically, these two pathways overlap in gall bladder where formation of biofilm by *S*. Typhimurium on gallstones plays an important role in protection from bile (38, 39). YciF had been identified as one of the downstream targets of CspE mediated bile stress response though the *yci*F deletion strain grows similar to WT in presence of bile (10). For pathogenic organism such as *S*. Typhimurium it is advantageous to have several proteins with overlapping substrate specificity and similar function to combat host defence and survive in hostile environments. An example is the *hpxF* deletion mutant wherein simultaneous deletion of five genes made the strain highly susceptible to oxidative stress (40). The bile stress response mediated by YciF is specific as Δ*csp*E is compromised in biofilm formation but overexpression of YciF could not rescue this phenotype (Fig. S7A and B). YciF is highly conserved across Enterobacteriaceae and the DUF892 domain is similar to ferritin. Ferritin forms a 24-mer cage like structure that can store the excess iron channelled into the mineralization sites of protein following the ferroxidase activity (41, 42). The oligomeric structure of Ferritin is critical for the storage of iron (13, 43). In this study we found that *S*. Typhimurium YciF is able to form higher order oligomeric complex which is of functional significance as it could allow the protein to store iron. YciF was found to efficiently bind and retain iron and the DUF892 domain was important for iron binding as it contains the putative metal binding sites. We also demonstrated that YciF has ferroxidase activity and, therefore, it possibly has ferritin-like function inside bacteria involving iron detoxification. Although YciF does not form cage like ferritin, its sequestration of excess iron within the helical backbone of oligomer may reduce accessibility of iron to dioxygen, hydrogen peroxide and superoxide resulting in a protective cellular function.

We also found that the *S*. Typhimurium Δ*csp*E strain displays disruption of expression of iron homeostasis genes when exposed to bile. This was confirmed by transcript level analysis of several genes required for uptake and utilization of iron. While the genes responsible for iron acquisition were significantly upregulated in Δ*csp*E compared to WT, expression of genes encoding proteins responsible for storage of excess iron such as *ftn*B and *dps* were downregulated. *dps* is a member of RpoS regulon, however, in *S*. Typhimurium sl1344, presence of *csp*C or *csp*E was shown to be required for optimal expression of *dps* (6, 11). Fluctuation in gene expression when encountering stress conditions is of adaptive importance to the pathogen (6). As bile can chelate iron (44) and act as a signal for iron restricted environment which the pathogen encounters inside host (45), activation of iron acquisition pathway along with repression of iron storage by *S*. Typhimurium WT possibly is a natural response to bile stress. However, the transcription profile of the Δ*csp*E strain indicates intracellular iron excess upon bile treatment, which can have deleterious effects by promoting ROS production.

Some studies suggest that there is an overlap between oxidative and bile stress response of *S*. Typhimurium. For example, the *sit*ABCD and manganese transport system *mnt*H is upregulated in both the stresses (24). In *E. coli*, bile salts have been shown to activate *mic*F and *osm*Y promoters which are associated with oxidative stress response (46). H_2_O_2_ is also generated endogenously by aerobic metabolism (40). Such correlations extend to oxidative stress and iron metabolism as well. *S*. Typhimurium challenged with H_2_O_2_ stress shows significant induction of iron acquisition proteins (47, 48). This could be as several proteins required to mitigate oxidative stress and associated DNA damage such as SodB and DNA glycosylase MutY require iron as cofactor (49–52). In this study, we were able to observe that presence of bile can cause a concurrent generation of ROS in *S*. Typhimurium, especially in Δ*csp*E. Consequently, the upregulation of iron acquisition pathway detected in WT strain could also be a response to oxidative stress generated due to bile treatment. In the Δ*csp*E strain though, that comprises of a cellular milieu with a markedly elevated ROS, the excessive induction of iron acquisition genes can cause toxicity through the Fenton reaction.

Among there are different reactive oxygen species, the Fenton reaction possibly is the most significant contributor to cell death owing to production of hydroxyl radicals which is considered the most potent reactive oxygen species (53–55). Even among the antibiotics, at least three major classes of bactericidal antibiotics, irrespective of their conventional targets have been found to stimulate formation of hydroxyl radicals which, in turn, cause bacterial killing. On the contrary, bacteriostatic antibiotics did not lead to production of hydroxyl radicals (54, 56). Subsequently, it was found that ROS accumulation is self-amplifying and once a threshold level of ROS is exceeded, cell death is sustained even after removal of initial stressor (57). Bile is also a bactericidal agent and in the Δ*csp*E strain causes impairment of iron balance. Pretreatment with iron chelator, 2,2 Bipyridyl significantly increased survival of Δ*csp*E during bile stress. This suggests that bile stress in Δ*csp*E strain causes cell lethality primarily through the Fenton reaction. Consequently, its inhibition by an iron chelator reduced the ROS generated in presence of bile and restored the growth similar to that of wild type (WT) strain. Reduction in levels of free iron can confer tolerance to various stress conditions that damage the cell through a common pathway of oxyradical generation. Similarly, the ferroxidase activity and subsequent chelation of iron by YciF overexpressed within the Δ*csp*E strain can contribute to its survival in presence of bile or peroxide stress by reducing the Fenton reaction (Fig 7). Unlike *S*. Typhimurium sl1344 where *csp*E and *csp*C are functionally redundant and the *csp*E*csp*C double deletion strain was susceptible to peroxide stress (11); we found that in *S*. Typhimurium 14028s, *csp*E deletion strain is sensitive to peroxide stress highlighting the strain-specific differences. No difference in growth with bile is found between the WT and Δ*yciF*, possibly due to redundant genes. However, overexpression of YciF significantly enhanced the survival of the *csp*E deletion strain in presence of peroxide. While Ferritin and Dps proteins are the general defence mechanisms to counter iron toxicity, our results indicate that YciF could complement this function in presence of physiologically relevant stressors. Overall, these results underscore the critical role of iron regulation in tolerance to bile mediated oxidative stress and peroxide stress in *S*. Typhimurium.

**Fig 7.**
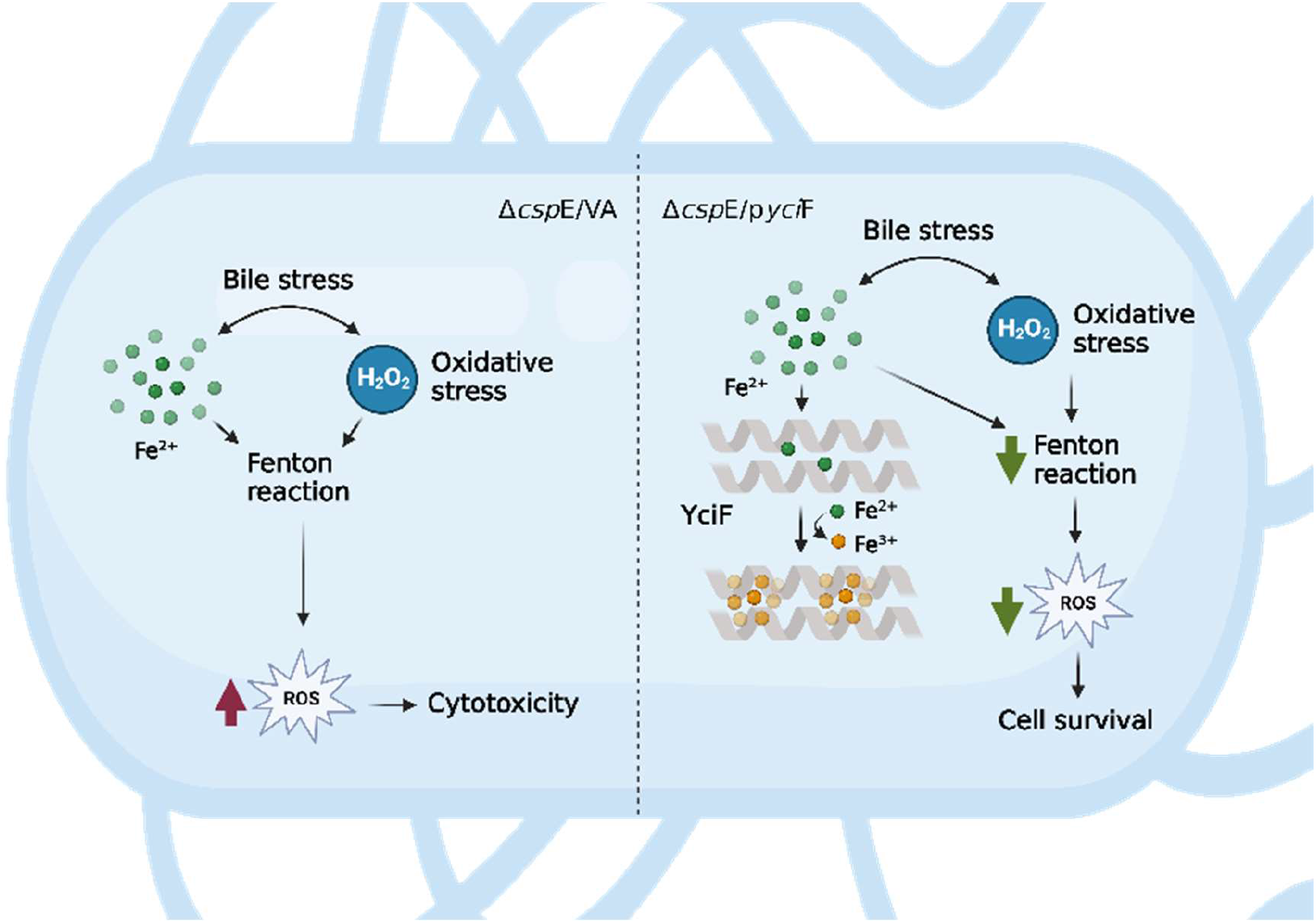
A model explaining the role of YciF during bile stress. Bile is known to cause an increase in oxidative stress. This, combined with H_2_O_2_ produced endogenously due to aerobic metabolism will undergo Fenton reaction, upon iron excess causing further sustained increase in ROS. Thus, the Δ*csp*E strain that shows disruption of iron homeostasis upon bile treatment, is likely to be more susceptible to Fenton reaction compared to the wild type strain. Higher ROS leads to cell death that can be mitigated by overexpression of YciF which binds to iron and has ferroxidase activity to help the Δ*csp*E strain detoxify the excess iron. Image was constructed using Biorender.com.

In this study we report for the first time, the physiological function of previously uncharacterized protein, YciF in bile stress response of enteric pathogen *S*. Typhimurium. YciF has ferroxidase activity that can prevent the damage to the cell caused by bile-mediated oxidative stress. The functional characterization of YciF can present a prototype for other homologous proteins of the DUF892 domain.

## Materials and Methods

### Bacterial strain and growth conditions

*S*. Typhimurium (ATCC 14028s) strain was used for in-vivo studies (10). Cultures (Table S1) were grown in Luria-Bertani (LB) broth at 37°C with aeration at 180 rpm. Overnight grown single colony culture was used as pre-inoculum for all experiments. When needed, Ampicillin was added at the concentration of 100 μg/mL. pQE60^ampR^ was used to overexpress yciF and its mutant forms.

### Multiple Sequence Alignment and phylogenetic tree construction

Homologous sequences closely related to *S*. Typhimurium (strain 14028s) YciF (Uniprot ID A0A0F6B221) were identified through UniprotKB database. Multiple sequence alignment was performed using Clustal omega program (58). Amino acid residues involved in metal ion coordination in the DUF892 domain were identified based on the YciF crystal structure (PDB ID 4ERU). For analysing wider taxonomic distribution, YciF sequence from *S*. Typhimurium (strain 14028s) was submitted to Jpred protein secondary structure prediction web server. Homologs of YciF from different genera were aligned to the input sequence and the results were shown using Jalview software (version 2.11.2.5). Phylogenetic tree was constructed using BLOSUM62neighbor joining method.

### Cloning yciF

PCR amplification of *yci*F was done using WT 14028s genomic DNA as template. The strains are listed in Table S2 and the primers used for site directed mutagenesis are listed in Table S3. PCR amplified *yci*F (with Strep II tag on C-terminal) and pQE60 plasmid (containing an IPTG inducible T5 promoter) were digested with BamHI and EcoRI (New England Biolabs) and purified by gel extraction (MinElute gel extraction kit, Qiagen, Germany). The purified insert was ligated into the cut pQE60 using T4 DNA ligase (Thermo Fisher Scientific) at 16°C overnight. p*yci*F construct was transformed into Top10 competent cells. p*yci*F was purified (GeneJET plasmid miniprep kit, Thermo Fisher Scientific, Lithuania) and confirmed by restriction digestion to visualize the insert release and Sanger sequencing (Aggrigenome, Kerala, India).

### Site directed mutagenesis

Site directed mutagenesis was performed using overlapping primers with desired mutations (Q54A, E113Q, E143D). 50 μL of PCR reaction mix was prepared using 100 ng template (pQE60 containing *yci*F), 250 nM forward and reverse primers, 0.2 mM dNTPs, 1X Phusion DNA polymerase buffer, 5% DMSO, 1 μL Phusion DNA polymerase. The negative control did not have DNA polymerase. Primer extension was done at 72 °C for 6 minutes (18 cycles). 1X Cutsmart buffer and 1 μL of Dpn1 (New England Biolabs) was added to 25 μL of PCR product and incubated at 37 °C for 4 hours to remove template DNA, followed by inactivation of Dpn1 at 80 °C for 20 minutes. Dpn1 digested product was used to transform Top10 competent cells by heat shock method. Plasmid was isolated from transformed cells and sequenced to confirm the mutation. *S*. Typhimurium WT and Δ*csp*E strains were transformed with pQE60 (vector alone), p*yci*F and p*yci*F containing Q54A, E113Q and E143D mutations by electroporation.

### Protein purification

Proteins were purified as described (59) with following modifications. *E.coli* BL21(DE3) containing either WT or mutant p*yci*F (Strep II tagged) was grown to OD_600_ of 0.5 at 37°C, 180 rpm in 1 L of LB supplemented with 100 μg/mL Ampicillin. The culture was induced with 1 mM IPTG (G biosciences, USA) followed by incubation at 20°C, 180 rpm for 12 hours. The cells were centrifuged at 6,000 rpm, 4°C for 15 minutes. Cell pellet was resuspended in lysis buffer (Buffer A) containing Tris-Cl pH 8, 10% glycerol, 150 mM NaCl and 1 mM DTT and sonicated. Lysate was centrifuged at 15,000 rpm, 4 °C for 45 minutes. The supernatant was subjected to affinity chromatography using 1 mL buffer equilibrated Strep-Tactin XT 4Flow resin (IBA Lifesciences-GmbH) and flow through was collected. Wash was given using buffer B (Tris-Cl pH 8, 10 % glycerol, 200 mM NaCl) and bound protein was eluted using buffer C (Tris-Cl pH 8.5, 10 % glycerol, 150 mM NaCl, 40 mM Biotin). Protein purity was confirmed on 12.5% SDS-PAGE gel with Coomassie brilliant blue staining and concentration of purified YciF was determined by Bradford assay. Protein was dialysed (in buffer containing Tris-Cl pH 7.5, 10% glycerol, 150 mM NaCl) and concentrated using Amicon ultra-15 centrifugal filter, 10 KDa cutoff (Merck-Millipore, Ireland) and stored at −80°C.

### Native PAGE

Purified proteins and native protein standard (Supelco-Merck) were loaded in 4-15% gradient mini protean precast native gel (Bio-Rad, USA). Electrophoresis was performed at 100 V, 4°C in 1X native running buffer. Gel was stained with Coomassie brilliant blue staining solution.

### Size exclusion chromatography

4 mg of native protein standard was loaded to calibrate the gel filtration column (Superdex 200 increase 10/300 GL column, GE Healthcare). Following this, 1 mg of purified protein (YciF, Q54A, E113Q and E143D) was loaded onto column equilibrated with elution buffer (Tris-Cl pH 8, 5% glycerol, 150 mM NaCl). All the runs were performed at 4°C using an AKTA purifier. Flow rate was maintained at 0.3 mL/min. Fractions were subjected to 12.5% SDS PAGE.

### Thermal shift assay

Thermal shift assay was performed as per Bio-Rad protocol with following modifications. 10 μg YciF was incubated with 200 μM (NH_4_)_2_Fe(SO_4_)_2_, MgSO_4_, ZnCl_2_ and EDTA in a hard-shell PCR 96 well plate (Bio-Rad, USA). 5X Sypro Orange dye (Supelco, Merck, USA) was added and change in fluorescence due to thermal denaturation was measured from 25°C to 95°C in a Bio-Rad CFX connect q-PCR instrument. To investigate thermal stability of WT and mutant proteins, Sypro Orange was added at 5X concentration to 10 μg of YciF, Q54A, E113Q and E143D and thermal denaturation was measured from 25°C to 95°C. Ramp rate was kept 0.5°C/10s. The first derivative of RFU was plotted against the temperature and the peak of first derivative was used to determine Tm.

### Iron staining

20 μM of purified proteins and BSA were incubated in absence or presence of 200 μM (NH_4_)_2_Fe(SO_4_)_2_ for 15 minutes on ice. The samples were loaded on 8% native gel and electrophoresis was performed at 75V, 4°C for 6 hours. Gel was stained with Ferene-S (Sigma-Aldrich, China) solution to detect bound iron (0.75 mM Ferene-S, 2% v/v acetic acid, 15 mM thioglycolic acid) in dark for 5 minutes (60). It was destained and stained again with Coomassie brilliant blue.

### Ferroxidase activity

Ferroxidase activity was determined as described previously (61–63) with following modifications. 10 mM (NH_4_)_2_Fe(SO_4_)_2_ stock solution was prepared in 0.1% (v/v) HCl. Buffer (25 mM HEPES, pH 7.3, 5% glycerol, 150 mM NaCl) was prepared anaerobically. 1 μM Protein was added to buffer and ferroxidase reaction was initiated by adding (NH_4_)_2_Fe(SO_4_)_2_ at different final concentrations in a UVMax 96-well plate (SPL life Sciences, Korea) with a final reaction volume of 200 μL. The absorbance was recorded at 310 nm after 600 s or every 30 s till 600 s using Infinite 200-Pro instrument (Tecan, Austria GmbH). For ferrous loss assay, Ferene-S was added at a final concentration of 500 μM after 600 s to the solution containing protein and 200 μM (NH_4_)_2_Fe(SO_4_)_2_ and absorbance was measured at 590 nm.

### RNA isolation

Bacterial cultures were grown to OD_600_ of 0.3 and then for 90 minutes in presence or absence of 3% bile. Cells were resuspended in 1 mL TRIzol reagent (Ambion, Invitrogen) and allowed to lyse for 30 minutes at 1500 rpm in a shaking drybath (DLAB). 250 μL of chloroform (Sigma-Aldrich, USA) was added and mixed vigorously for 30 s. The sample was allowed to stand for 2 minutes and centrifuged at 12,000 x g, 4 °C, 15 minutes. The aqueous phase was mixed with 200 μL of chloroform and phase separation was performed again as earlier. 300 μL of aqueous phase was added to 500 μL of 2-propanol (Merck, Germany) and kept overnight at −20°C for RNA precipitation. The RNA was pelleted at 15,000 x g, 4 °C, 30 minutes and washed with 75% ethanol twice. Pellet was kept for drying to remove residual ethanol and dissolved in nuclease-free water. RNA integrity was analysed by electrophoresis on a 1.5% agarose gel. Concentration was determined using Nano-Drop (Thermo Fisher Scientific).

### qRT-PCR

qRT-PCR was performed as described previously (10) with following modifications. Genomic DNA was removed from the extracted RNA using DNase I (New England Biolabs) treatment and removal was confirmed by PCR of a reference gene. 2.5 μg of DNase I treated RNA was reverse transcribed using random hexamer and RevertAid cDNA synthesis kit (Thermo Fisher Scientific, Lithuania). q-PCR was performed using 100 ng of cDNA sample, 250 nM of gene specific primer, SYBR green master mix (BioRad) in a Bio Rad CFX connect q-PCR instrument. The primers (Sigma-Aldrich, Bangalore) for q-PCR are listed in Table S4. The relative quantities of transcripts were estimated using ΔΔCq method normalized against two reference genes, *gm*k and *gyr*B. The WT strain grown in LB was used as control sample.

### Bacterial stress assays

Experiments were done in 5 mL LB. Overnight grown cultures of indicated *S*. Typhimurium WT and Δ*csp*E strains were normalized to OD_600_ of 2, diluted 1:250 in LB and grown to OD_600_of 0.2, followed by addition of stress inducing compounds. Cells were grown in 50 mL falcons for 6 hours (3 hours in H_2_O_2_), at 180 rpm, 37°C. 200 μL of culture from each growth condition was taken in clear flat bottom 96-well plate and OD_600_ was measured using Infinite 200-Pro instrument (Tecan, Austria GmbH).

### ROS estimation

ROS estimation was done as described previously (64). *S*. Typhimurium WT and Δ*csp*E strains were diluted 1:250 in LB and grown to OD_600_ of 0.3 and treated with 3% bile for 6 hours. Cells were centrifuged and washed with 1X PBS. Cells were incubated with 20 μM 2’,7’-Dichlorofluorescin Diacetate (DCFDA) (Millipore, China) at 37°C for 30 minutes. Cells were washed twice with 1X PBS and immediately analyzed by flow cytometry. 20,000 cells were recorded per condition in a FACS Verse instrument (BD Biosciences, USA). Data was analyzed using FlowJo software (version 10). For quantitative estimation of ROS, 200 μL of cell suspension from each condition was taken in 96-well plate and fluorescence was measured at excitation of 485 nm and emission of 535 nm using Infinite 200-Pro instrument (Tecan, Austria GmbH). The values were normalized to OD_600_.

### Bile tolerance assay in presence of 2,2-bipyridyl

*S*. Typhimurium WT and Δ*csp*E strains were diluted 1:250 in LB with or without 200 μM 2,2-bipyridyl (Sigma-Aldrich, China). Cultures were incubated at 37°C at 180 rpm, grown to OD_600_ of 0.2 and then treated with bile (Sigma-Aldrich) at a final concentration of 3% for 6 hours. OD_600_ was measured and cells were plated on LB agar at appropriate dilutions to count CFUs. Qualitatively, viability post bile treatment was determined by spotting serial dilutions on LB agar. CFU plates were kept at 37°C while spotting plates were kept at 30°C overnight to prevent overgrowth.

### Statistical analysis: Data were analyzed using GraphPad prism software

Two-way ANOVA was used to analyze bacterial growth and ROS estimation with each strain and condition compared using Sidak’s multiple comparisons test. One way ANOVA was used to analyze q-PCR data.

## Acknowledgements

We thank the SEC-MALS facility at the Molecular Biophysics Unit, Indian Institute of Science and FACS facility, Division of Biological Sciences, Indian Institute of Science. We appreciate the help of Smruti Nayak and Pragya Ahuja with the SEC, Kshitiza Mohan Dhyani and Sirisha Jagdish with protein purification and Siddharth Jhunjhunwala for Biorender accession.

