## Supplemental Information for "Functional characterization of *Salmonella* Typhimurium encoded YciF, a domain of unknown function (DUF892) family protein, and its role in protection during bile and oxidative stress"

**Supplementary Information:**

| **Table S1**: List of strains and plasmids Strain | Genotype/Features | Source/Reference |
|---|---|---|
| *S*. Typhimurium ATCC 14028s | Wild type, parent of all *Salmonella* strains | Ray *et al*., 2019 (Ref 10 in main manuscript) |
| *E.coli*TOP10 | F– *mcr*A Δ(*mrr-hsd*RMS-*mcr*BC) φ80*lac*ZΔM15 Δ*lac*X74 *rec*A1 *ara*D139 Δ(*ara-leu*)7697 *gal*U*gal*K λ– *rps*L(StrR) *end*A1 *nup*G | ThermoFisher Scientific |
| *E.coli*BL21(DE3) | F– *omp*T*hsd*SB (rB–, mB–) *gal dcm*(DE3) | Ray *et al*., 2019 |
| Δ*csp*E | 14028s; *csp*E::FRT | Ray *et al*., 2019 |
| WT/VA | 14028s; pQE60; $Amp^r$ | This study |
| Δ*csp*E/VA | 14028s; pQE60; $Amp^r$ | This study |
| WT/p*yci*F | 14028s; pQE60; $Amp^r$ | This study |
| Δ*csp*E/p*yci*F | 14028s; pQE60; $Amp^r$ | This study |
| WT/p*yci*F-Q54A | 14028s; pQE60; $Amp^r$ | This study |
| Δ*csp*E/ p*yci*F-Q54A | 14028s; pQE60; $Amp^r$ | This study |
| WT/p*yci*F-E113Q | 14028s; pQE60; $Amp^r$ | This study |
| Δ*csp*E/p*yci*F-E113Q | 14028s; pQE60; $Amp^r$ | This study |
| WT/p*yci*F-E143D | 14028s; pQE60; $Amp^r$ | This study |
| Δ*csp*E/p*yci*F-E143D | 14028s; pQE60; $Amp^r$ | This study |
| WT/VA | 14028s; pRS424; $Amp^r$ | Ray *et al*., 2019 |
| Δ*csp*E/VA | 14028s; pRS424; $Amp^r$ | Ray *et al*., 2019 |
| WT/p*csp*E | 14028s; pRS424; $Amp^r$ | Ray *et al*., 2019 |
| Δ*csp*E/p*csp*E | 14028s; pRS424; $Amp^r$ | Ray et al., 2019 |

| **Table S2**: Plasmids | Genotype/Features | Source/Reference |
|---|---|---|
| pQE60 | A 3431 bp bacterial expression plasmid. Contains T5 promoter and confers ampicillin resistance | Qiagen |
| p*yci*F | pQE60 with *yci*F cloned between *Bam*HI and *Eco*RI | This study |

| | | |
|---|---|---|
| p*yci*F-Q54A | pQE60 with Q54A mutant of *yci*F cloned between *Bam*HI and *Eco*RI | This study |
| p*yci*F-E113Q | pQE60 with E113Q mutant of *yci*F cloned between *Bam*HI and *Eco*RI | This study |
| p*yci*F-E143D | pQE60 with E143D mutant of *yci*F cloned between *Bam*HI and *Eco*RI | This study |

| **Table S3**: Primers for cloning and site directed mutagenesis | Sequence 5'-3' |
|---|---|
| *yci*F cloning FP | CGCGAATTCATGAATATCAAAACCGTTGAAG |
| *yci*F cloning RP | CGCGGATCCTTATTTTTCGAACTGCGGGTGGCTCCACGCGCTTTTCGATTTGCGTTCAGCAC |
| Q54A FP | AAGAAACCCAGGGTGCGATTGAACGTATTG |
| Q54A RP | CAATACGTTCAATCGCACCCTGGGTTTCTT |
| E113Q FP | AGTCGAGCATTACCAAATCGCCAGCTA |
| E113Q RP | TAGCTGGCGATTTGGTAATGCTCGACT |
| E143D FP | CCCTCGACGAGGATAAACAAACTGATT |
| E143D RP | AATCAGTTTGTTTATCCTCGTCGAGGG |

| **Table S4**: Primers used for q- PCR | Sequence 5'-3' |
|---|---|
| *feo*B FP | TCCGTCGCCATCTTTATTCA |
| *feo*B RP | GTACATCATCCCGATTTGCG |
| *fep*A FP | TGGGTAACGATGATCTGAAA |
| *fep*A RP | TCTCCCACTGATAAACATCC |
| *fhu*C FP | TGCATCGTTTAAGCCAACAG |
| *fhu*C RP | CGTAAATCTGTTCCAGCGTG |
| *sit*A FP | GTGCGTAAAGTGATTGATACCA |
| *sit*A RP | ATCCAGATAGGTTGGCACAG |
| *fes* FP | AAAGACCGTTGGCTATTCTG |
| *fes* RP | GTGTTGAGTATCAATCGCGT |
| *fhu*F FP | ATGGGCGCAATGGTATATCG |
| *fhu*F RP | GCACATCAAGCCAGAAACAC |
| *yqj*H FP | GGCGCTGGATTTCTTTATCC |

| | |
|---|---|
| *yqj*H RP | TCGCAGACATAAACCTGACA |
| *ftn*B FP | GCTCTGCTCTCTGGAAGAAT |
| *ftn*B RP | TAAAAGTACGCCATCCTGCT |
| *dps* FP | TCCACTGAAAAGCTATCCGC |
| *dps* RP | GTGATGCGGCGGTAAAGATA |
| *fur* FP | ATCACGACCATCTTATCTGC |
| *fur* RP | GGCTGTGATTAGTTAAACGA |

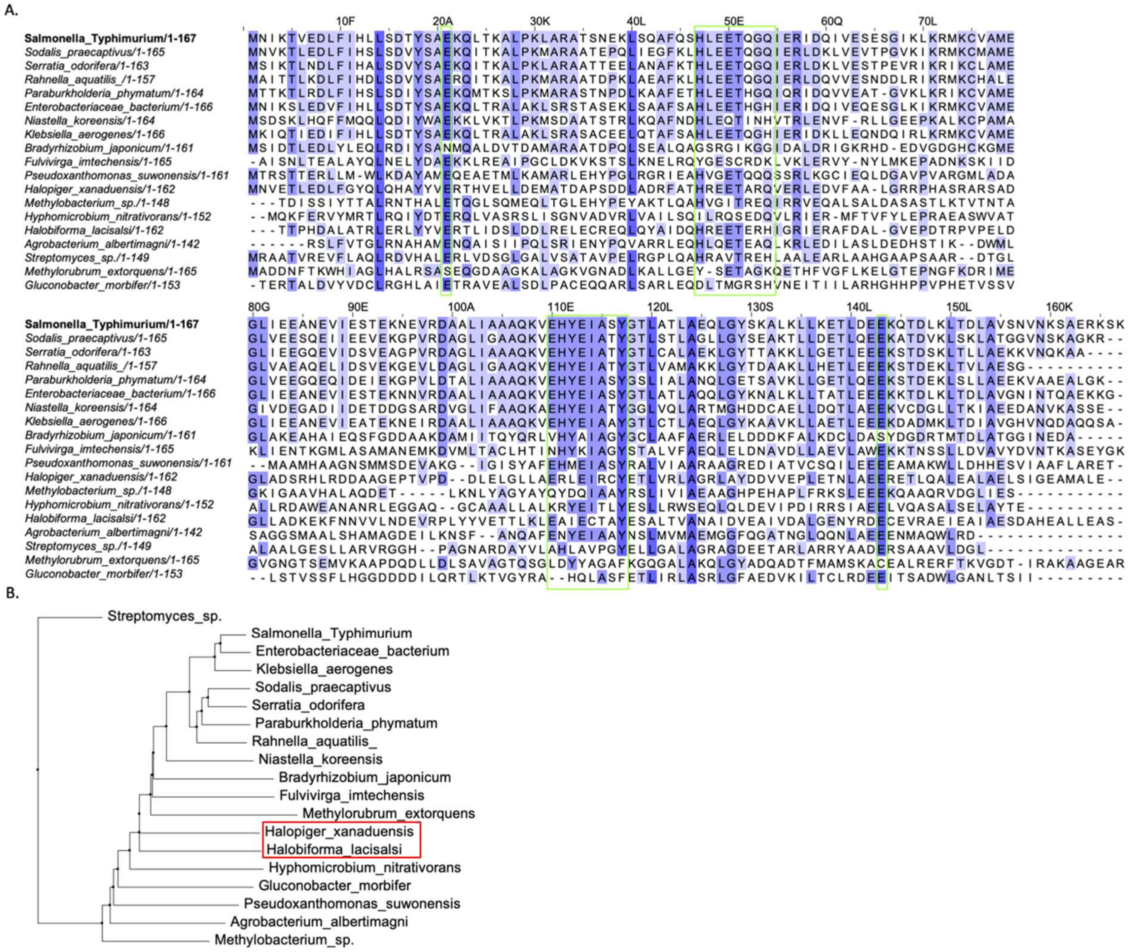

**Fig S1.** Taxonomic distribution of the DUF892 domain. (A) Multiple sequence alignment was carried out using *S*. Typhimurium YciF as primary sequence. The intensity of blue colour corresponds to percentage sequence identity of residues across different genera with deep blue representing high sequence identity. Metal binding motifs and coordination residues are boxed in green. (B) Phylogenetic tree representing evolutionary distances of *S*. Typhimurium YciF and its DUF892 domain homologs across distinct taxonomic groups. Members of Archaea are boxed in red.

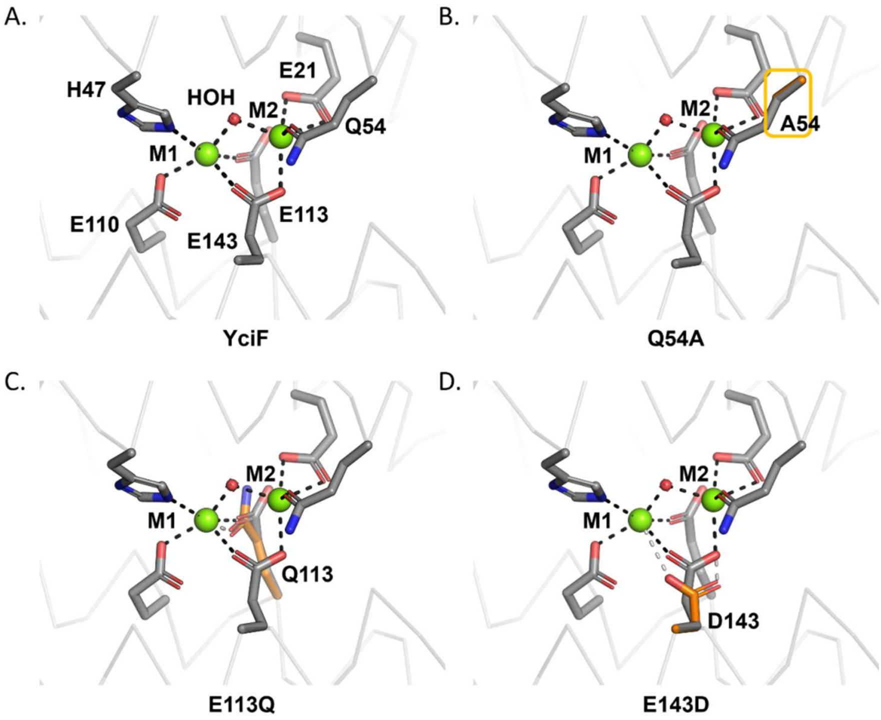


**Fig S2.** In-silico comparison of metal binding sites (M1 and M2) of YciF and the site-specific mutants. (A) *S*. Typhimurium YciF crystal structure has Glu21, Gln54 coordinating M2 site whileGlu110, Glu113 coordinate M1 site. Glu143 and water molecule act as bridging ligands for both the metal binding sites. (B) Substitution of Gln54 with alanine results in incomplete coordination at the M2 site. Rectangle marks the position of mutated residue. (C) The coordination remains unchanged in negatively charged Glu113 to uncharged glutamine substitution. (D) Substitution of bidentate Glu143 with aspartate creates an increased coordination distance to both the metal ion and coordinating residue thereby weakening it. Figures were prepared using PyMol v2.4.

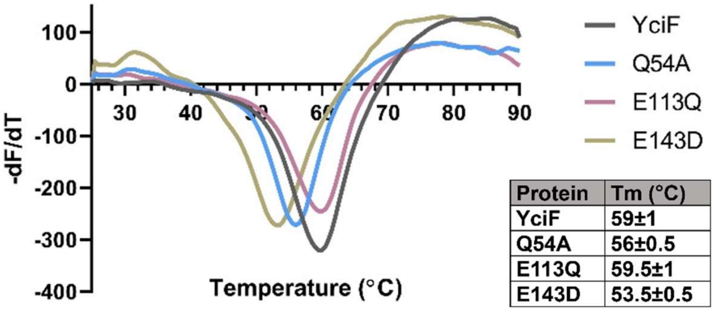


| Protein | Tm (°C) |
|---|---|
| YciF | 59±1 |
| Q54A | 56±0.5 |
| E113Q | 59.5±1 |
| E143D | 53.5±0.5 |

**Fig S3.** Thermal shift assay to determine the stability of *S*. Typhimurium YciF and DUF892 mutants. Sypro Orange was added at 5X concentration to 10 µg of YciF, Q54A, E113Q and E143D and thermal denaturation was recorded from 25°C to 95°C at a ramp rate of 0.5°C/10s. Tm corresponds to temperature at the derivative peak. The data is representative of 4 independent experiments.

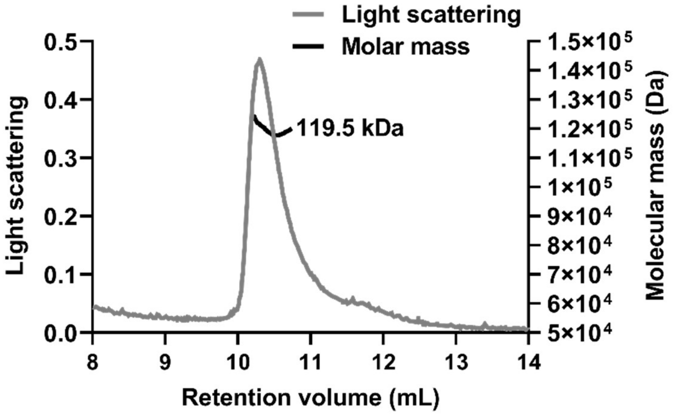


**Fig S4.** Molar mass of purified YciF as determined by SEC-MALS. Purified YciF was passed through Superdex S-200 column. The chromatogram displays light scattering at 90° angle along with molar mass of the peak determined by MALS.

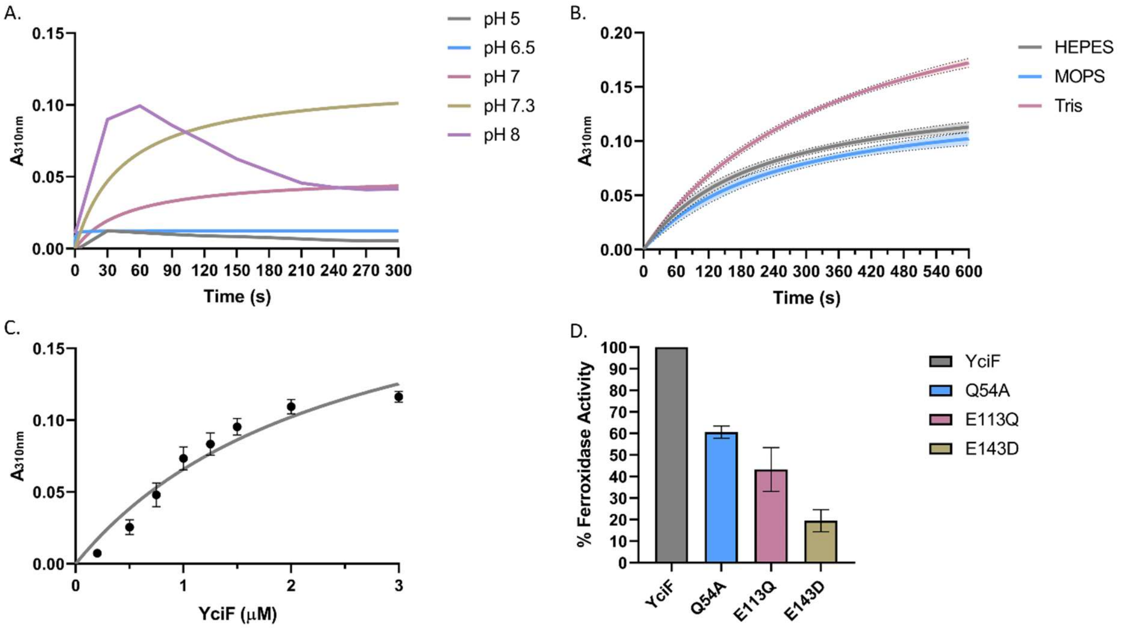


**Fig S5.** (A) Optimization of pH for ferroxidase assay. 200 µM of $(NH_4)_2Fe(SO_4)_2)$ was added to 1 µM of purified YciF and absorbance at 310 nm measured for 300s at 30 s interval. (B) Buffer condition standardization for ferroxidase assay. 200 µM of $(NH_4)_2Fe(SO_4)_2)$ was added to 1 µM of purified YciF in HEPES, MOPS and Tris buffer (pH 7.3) and absorbance at 310 nm measured for 600s at 30 s interval. (C) Ferroxidase activityrecorded in presence of different concentrations of purified YciF and 200 µM of $(NH_4)_2Fe(SO_4)_2)$. Absorbance was measured at 310 nm after 600s. (D) Ferroxidase activity of YciF and the mutants were compared by the ferrous loss assay using Ferene-S, a chromogen that forms prussian blue coloured complex with $Fe^{2+}$. 1µM of purified YciF, Q54A, E113Q and E143D were incubated with200 µM of $(NH_4)_2Fe(SO_4)_2)$ at 25°C for 15 minutes. Ferene-S solution was added, and absorbance was measured at 590 nm.The data is representative of 3 independent experiments.

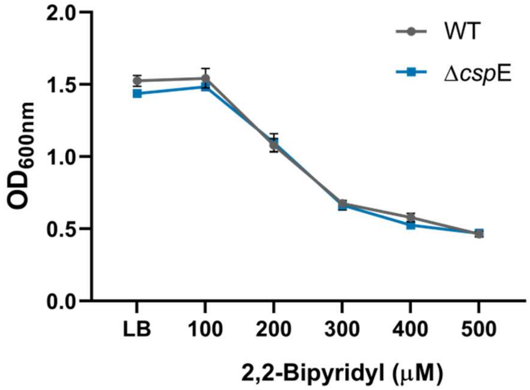


**Fig S6.** *S.*Typhimurium WT and Δ*csp*E strains have similar growth in presence of iron chelator, 2,2-Bipyridyl. Strains were grown in different concentrations of Bipyridyl for 6 hours and O.D. was measured at 600 nm. Data shown as mean±SD and is representative of 2 independent experiments.

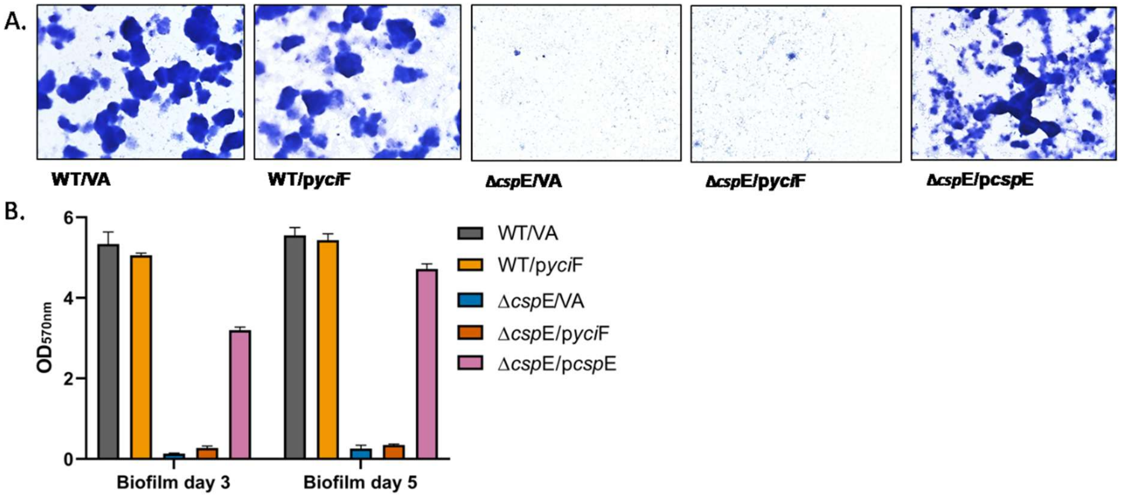


**Fig S7.** The Δ*csp*E strain expressing *yci*F is compromised in biofilm formation. (A) Indicated *Salmonella* strains were added to LB-NaCl in a 12-well plate. Δ*csp*E/pQE60 was used as vector alone control while Δ*csp*E/p*csp*E was used as positive control. Plate was kept at 28°C for 3 and 5 days to allow biofilm formation. Biofilms were stained with 1% crystal violet staining solution and visualized under microscope. (B) Biofilm formation was quantitated using 33% acetic acid as destaining solution and measuring O.D. at 570 nm. The data is representative of 3 independent experiments.

**Supplementary Material and Methods:**

**SEC-MALS:** SEC-MALS was done as described (i). Briefly, 200 µg protein was injected into the analytical gel filtration column (Superdex 200 increase 10/300 GL column, GE Healthcare) equilibrated with elution buffer (Tris-Cl pH 8, 5 % glycerol, 150 mM NaCl). The in-line UV system (Shimadzu) was coupled to the MALS system. MALS signal was detected by miniDAWN TREOS MALS detector (Wyatt Technologies orp, USA) and a refractive index detector (Waters 24614). The UV, RI and MALS data was analysed using ASTRA (version 6.1) software.

**Biofilm formation:** Biofilm formation was performed as described (ii). Overnight grown *S*. Typhimurium wild type and Δ*csp*E strains were normalized to $OD_{600}$ of 3. Fifty µL of normalized culture was added to 1.5 mL of LB without NaCl in a 12-well tissue culture grade sterile microtitre plate (Tarsons, Korea). The plate was covered and kept in 28 °C under stagnant condition. After day three and day five, planktonic cells were removed, and biofilms were washed twice with 1X PBS to remove unadhered cells. Biofilms were dried and heat fixed at 60 °C for 1 hour. Crystal violet staining solution (0.33%) was used to stain biofilm at room temperature for 10 minutes. The excess stain was removed by washing with 1X PBS, thrice. Images for biofilms were acquired at 20X magnification. Acetic acid (33%) was used as destaining solution to extract stain and absorbance was measured at 570 nm.

ii) Ray S, Da Costa R, Thakur S, Nandi D. 2020. Salmonella typhimurium encoded cold shock protein e is essential for motility and biofilm formation. *Microbiol* (United Kingdom), 10.1099/mic.0.000900.
